# CDK activity at the centrosome regulates the cell cycle

**DOI:** 10.1101/2023.08.16.553560

**Authors:** Emma L. Roberts, Nitin Kapadia, Tania Auchynnikava, Souradeep Basu, Paul Nurse

## Abstract

Cyclin-dependent kinases (CDKs) complexed with cyclins drive progression through the eukaryotic cell cycle. From yeast to human cells, cyclin-CDK localises to the centrosome, but the importance of this localisation is unclear. The conserved ‘hydrophobic patch’ substrate docking site on human Cyclin B1 and on the equivalent fission yeast Cdc13 mediates their localisation to the centrosome and the spindle-pole body (SPB, yeast centrosome equivalent). A hydrophobic patch mutant (HPM) of Cdc13 cannot enter mitosis, but whether this mitotic defect is due to defective SPB localisation or defective cyclin-substrate docking is unknown. Here we show that artificially restoring Cdc13^HPM^ SPB localisation in fission yeast partially rescues both mitosis and defective CDK substrate phosphorylation at both the SPB and within the cytoplasm. In addition, we found that an HPM of the S-phase cyclin Cig2 has defective SPB localisation but is still able to perform bulk DNA synthesis. Our results demonstrate that the hydrophobic patch mediates the SPB localisation of both S- and M-phase cyclins, and that Cdc13 SPB localisation is essential for mitotic entry and for full phosphorylation of CDK substrates, supporting the view that the centrosome plays a role as a signalling hub regulating CDK cell cycle control.

## Introduction

Cyclin-CDK complexes are the master regulators of the eukaryotic cell cycle, phosphorylating hundreds of substrates to orchestrate critical events including DNA replication and mitosis. Cyclin-CDK activity is regulated in multiple ways, including through direct docking to substrates and the dynamics of its cellular localisation, both of which involve a conserved site on cyclins known as the hydrophobic patch^1–7^. From yeast to human cells, several core cell cycle regulators including cyclin-CDK localise to the centrosome^5,6,8–11^, which has been suggested to act as a signalling centre^12,13^. For example, signalling at the centrosome in the early embryo of *C. elegans* can influence the timing of mitotic entry as assayed by nuclear envelope breakdown^14,15^, suggesting that centrosomes may act as a signalling hub regulating mitotic entry. Supporting this, during mitosis an active form of Cyclin B1-Cdk1 is first detected at the centrosomes in HeLa cells^16^. However, centrosomes are not required for cell cycle progression or cell division in *Drosophila*^17^, and removal of centrosomes in some cultured mammalian cell lines does not prevent mitotic entry or progression^12,13,18,19^. Therefore, although cyclin-CDK has been seen to localise to the centrosome in many eukaryotic species, questions remain around how important this conserved phenomenon is for cell cycle regulation.

The relative simplicity of the cell cycle in the fission yeast *Schizosaccharomyces pombe*, in which the mitotic cyclin-CDK complex Cdc13-Cdc2 is able to drive the whole cell cycle when all other cell cycle cyclins are genetically deleted, makes it an advantageous system for unravelling complex issues involved in CDK regulation of the cell cycle^20^. Work on the fission yeast spindle-pole body (SPB, the yeast centrosome equivalent) has established that signalling based around the SPB component Cut12 can influence mitotic entry through the mitotic regulator Plo1 (Polo kinase, known as Plk1 in humans)^21–24^. Polo kinase contributes to feedback loops that act on an inhibitory phosphorylation of CDK residue Y15, a major point of CDK activity regulation, and removal of Y15 phosphorylation promotes mitotic entry^25–34^. Thus, Plo1 regulation at the SPB is thought to promote feedback activity on CDK to regulate mitotic entry^24^. Furthermore, artificial recruitment of Cdc2^Y15F^ (which is insensitive to the inhibitory protein kinase Wee1) to the SPB is sufficient to promote mitotic entry in G2 cells^35^. However, it is not known if cyclin-CDK localisation to the SPB is needed for mitosis.

We have previously shown that the hydrophobic patch docking site on cyclins, which is needed for the centrosomal localisation of mammalian Cyclin A2^6^, is also involved in the localisation of the mitotic cyclin Cdc13 to the SPB in fission yeast and the centrosomal localisation of human Cyclin B1^7^. A hydrophobic patch mutant (HPM) of Cdc13, in which three residues previously shown to be critical to hydrophobic patch function are mutated to alanine^2^, does not support cell viability as the hydrophobic patch is needed for mitosis^7^. However, in addition to correct localisation of cyclin-CDK, the hydrophobic patch is also known to dock to CDK substrates and regulators, enhancing substrate phosphorylation or influencing CDK activity regulation, and these are disrupted upon mutating the hydrophobic patch^1–3,36,37^. Thus, it remains unknown whether the essential *in vivo* function of the Cdc13 hydrophobic patch lies in its role in cyclin-CDK SPB localisation, or its role in substrate docking. It is also unclear to what extent the different *in vivo* functions of the cyclin hydrophobic patch are conserved between S- and M-phase cyclins. The hydrophobic patch of budding yeast S- and M-phase cyclins enhance *in vitro* phosphorylation of different substrates^3^, and can recognise different docking motifs on substrates^36,38^, indicating some differences in the hydrophobic patch of different cyclins. However, the hydrophobic patch of both mammalian Cyclins A2 and B1 mediate their centrosomal localisation^5–7^.

Here we investigate the importance of the hydrophobic patch and centrosomal cyclin-CDK localisation in S- and M-phase cyclins *in vivo* using the relatively simple *S. pombe* cell cycle. Our results demonstrate that SPB localisation of the mitotic cyclin Cdc13 is essential for mitotic entry and that the hydrophobic patch of the S-phase cyclin Cig2 is not essential for bulk DNA replication but is involved in SPB localisation.

## Results

### Cells driven by Cdc13^HPM^ arrest in G2 without Cdc13^HPM^ SPB localisation

We first investigated whether cells expressing only Cdc13^HPM^ arrest in the cell cycle without undergoing Cdc13^HPM^ SPB localisation. We previously found that, in the presence of wild-type Cdc13, an exogenous copy of Cdc13^HPM^-sfGFP (HPM: M235A, L239A, W242A) does not accumulate at the SPB in early G2, but is able to do so in late G2/early mitosis^7^. However, in the absence of wild-type Cdc13 when cells only have Cdc13^HPM^, they arrest in G2^7^. Therefore, we have hypothesised that cells expressing only Cdc13^HPM^ arrest before the point in the cell cycle at which Cdc13^HPM^ accumulates at the SPB. To test this, we introduced into cells an exogenous copy of either Cdc13^WT^-sfGFP^39^ or Cdc13^HPM^-sfGFP and placed wild-type Cdc13 under a thiamine-repressible promoter (Fig. 1a). In addition, the G1/S cyclin genes *cig1* and *cig2* were deleted and an SPB component, Sid4-mRFP^40^, was used as an SPB marker. Cells were followed after release from a G1 arrest in the presence of thiamine to repress wild-type *cdc13* (Fig. 1b). As expected from our previous findings^7^, cells expressing Cdc13^WT^-sfGFP and Cdc13^HPM^-sfGFP both performed S-phase with similar timings (Fig. 1c). However, although cells expressing Cdc13^WT^-sfGFP underwent nuclear division to form binucleates, indicating progression through mitosis, cells expressing Cdc13^HPM^-sfGFP were unable to do so (Fig. 1d).

**Figure 1:**
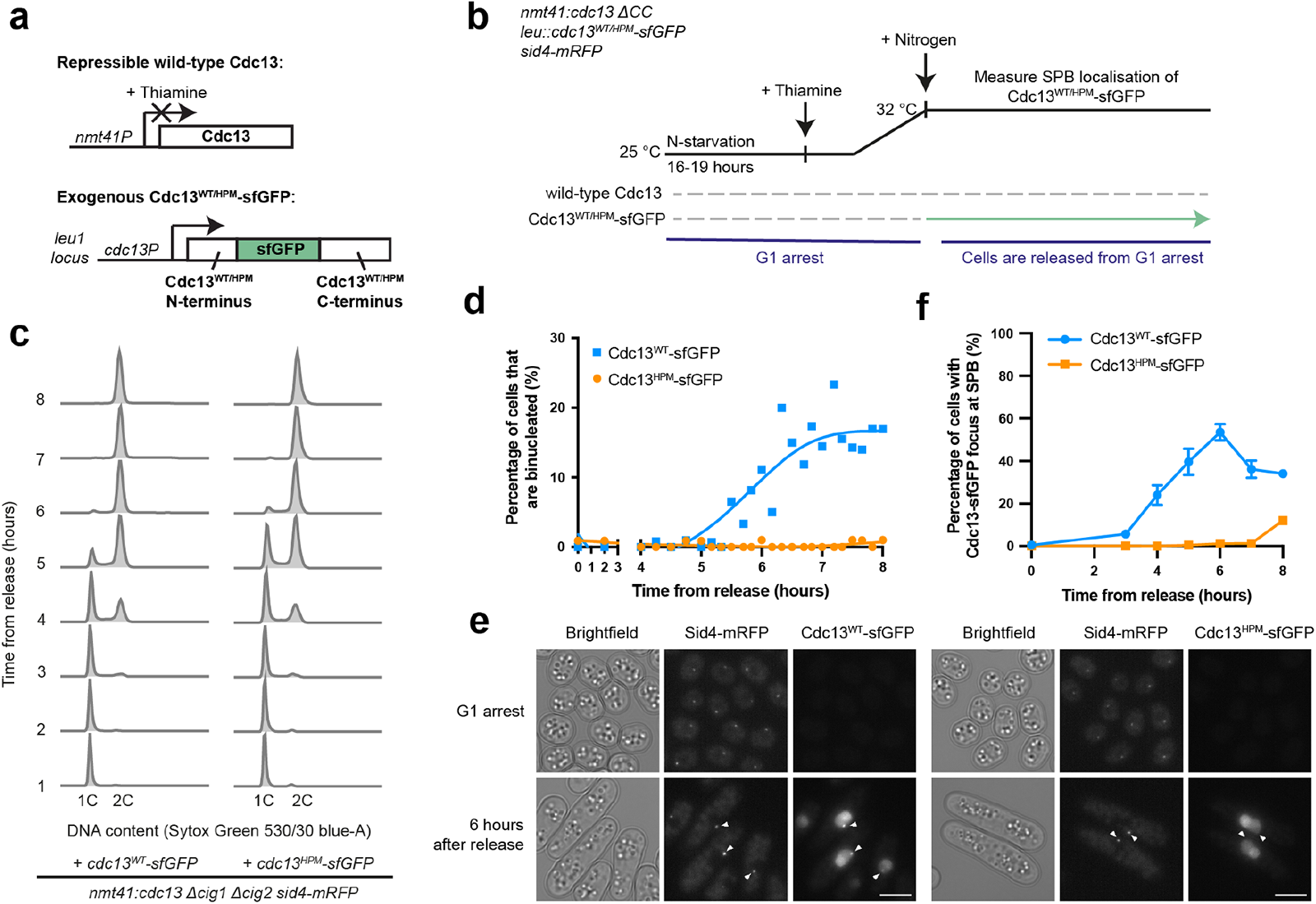
Cells driven by Cdc13^HPM^ arrest in G2 without detectable Cdc13^HPM^ SPB enrichment. **a** Schematic of Cdc13 conditional expression system. Wild-type Cdc13 is expressed under the thiamine-repressible *nmt41* promoter. Cells also contain an exogenous copy of either *cdc13^WT^-sfGFP* or *cdc13^HPM^-sfGFP*, expressed under the *cdc13* promoter. The sfGFP tag is internal to Cdc13^39^. **b** Experimental design. Cells were starved of nitrogen overnight resulting in a G1 arrest, during which Cdc13 protein is degraded. Thiamine was added to the culture one hour before release, and maintained throughout the experiment, to repress the transcription of wild-type *cdc13*. Cells were released from the G1 arrest by addition of nitrogen, and samples were taken to assay cell cycle progression and exogenous Cdc13-sfGFP localisation. **c** DNA content of cells after release from the G1 arrest, determined by flow cytometry. **d** Heat-fixed cells were stained with DAPI and scored for binucleation to measure passage through mitosis. n > 70 cells per time point, spline was plotted in GraphPad Prism. **e** Images showing Cdc13-sfGFP accumulation and localisation in cells arrested in G1 and 6 hours after release. Fluorescence images are maximum projection images, and the same pixel range has been applied to all images of the same channel. Arrows indicate location of SPBs, judged by Sid4-mRFP signal. Scale bars, 5 μm. **f** Cells were scored for a focus of Cdc13-sfGFP colocalised with the SPB marker Sid4-mRFP. Mean and SD of two biological replicates are shown, apart from time-points 3 (WT) and 8 (WT and HPM), where there is data from one replicate. n > 240 cells per strain per timepoint.

Using an automated spot detection pipeline (Supplementary Fig. 1) we investigated the SPB localisation of Cdc13^WT^-sfGFP and Cdc13^HPM^-sfGFP. The proportion of cells with Cdc13^WT^-sfGFP at the SPB increased at 3 hours after release from G1 arrest, peaking at roughly 50% of cells after 6 hours, and then decreased as cells underwent mitosis and degraded Cdc13 (Fig. 1d-f). In contrast, the proportion of cells with Cdc13^HPM^-sfGFP at the SPB remained negligible after release (Fig. 1e,f). We conclude that the vast majority of cells driven by Cdc13^HPM^-sfGFP arrest in G2 without detectable Cdc13^HPM^-sfGFP present at the SPB.

### Artificial restoration of Cdc2 SPB localisation

The G2 arrest of cells driven by Cdc13^HPM^ could be caused by the lack of Cdc13^HPM^ SPB localisation, or a disruption of cyclin-substrate docking, or both. To test the importance of Cdc13 SPB localisation, we investigated whether artificially restoring Cdc13^HPM^ SPB localisation rescued the mitotic defect of Cdc13^HPM^. To do this, we adapted a system previously used to tether CDK to the SPB, which demonstrated that CDK signalling at the SPB component Cut12 can regulate mitosis^35^ (Supplementary Fig. 2a-c). We ensured that Cdc13^HPM^ was the only copy of Cdc13 expressed during the experiment (Supplementary Fig. 2a), and we tethered an exogenous copy of the analogue-sensitive allele Cdc2(as)-M17-GBP-mCherry^41^ (sensitive to inhibition by 1-NmPP1) to Cut12-NEGFP^42,43^ (Supplementary Fig. 2c).

This experiment aimed to determine whether activating the SPB-tethered Cdc2(as)-M17-GBP-mCherry would allow cells to progress past the G2 arrest caused by Cdc13^HPM^. We first arrested cells in G1, repressing wild-type *cdc13* expression as well as inducing Cdc2(as)-M17-GBP-mCherry expression but in the presence of 1-NmPP1 to inhibit its CDK activity (Fig. 2a,b). Thus, the only active copy of Cdc2 present in cells was the endogenous Cdc2. Upon release from G1 arrest, cells performed S-phase but arrested in G2 as expected for cells driven by Cdc13^HPM^ (Supplementary Fig. 2d,e). Once cells had accumulated in G2, 1-NmPP1 was removed from the cultures to activate Cdc2(as)-M17-GBP-mCherry, and cells were monitored for progression through mitosis (Fig. 2a).

**Fig 2:**
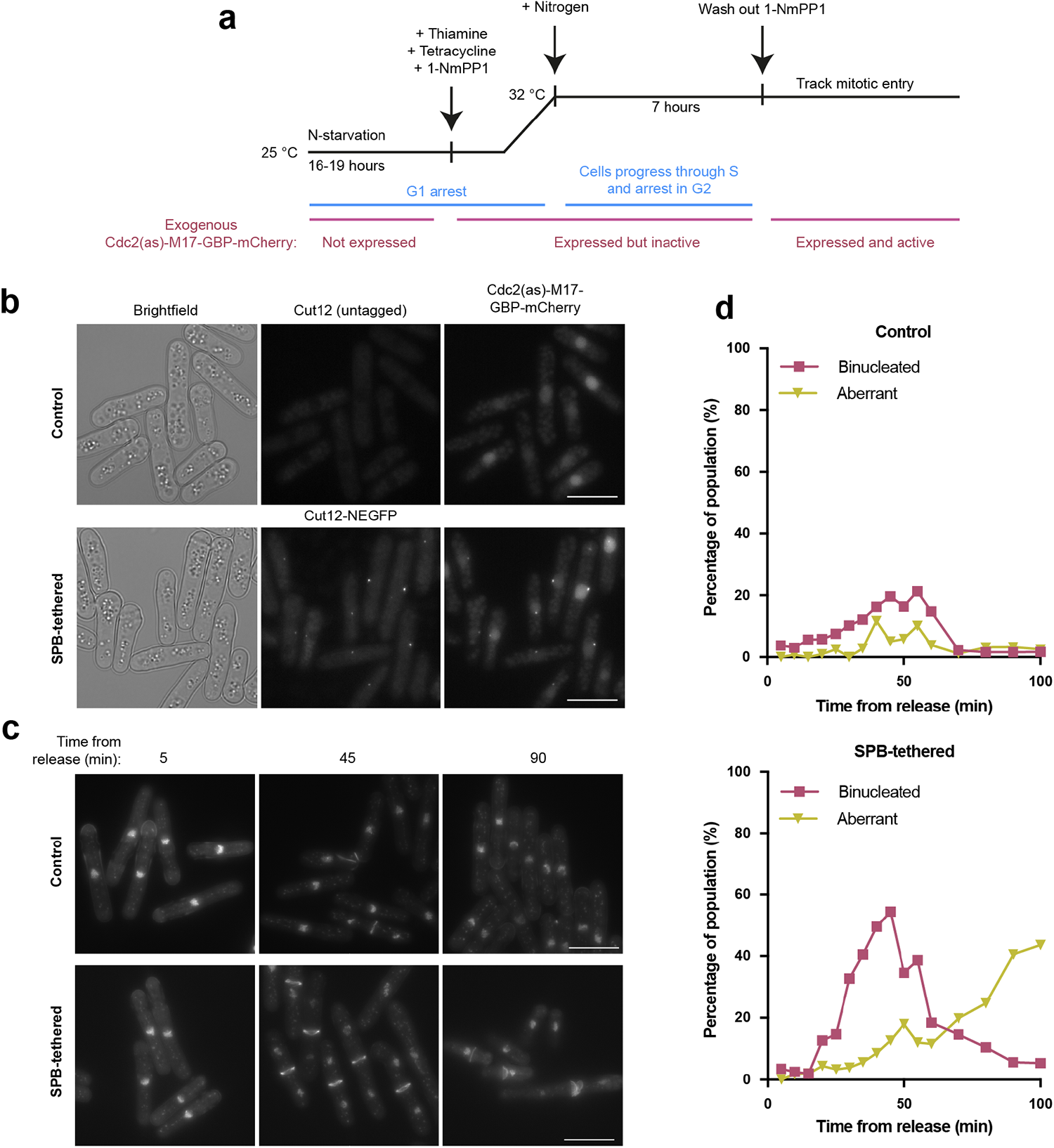
Artificially restoring Cdc13^HPM^ SPB localisation promotes mitotic entry. **a** Experimental outline. Cells were starved of nitrogen for between 16-19 hours to arrest them in G1. Thiamine (to repress wild-type Cdc13 transcription), tetracycline (to induce Cdc2(as)-M17-GBP-mCherry expression) and 1-NmPP1 (to inhibit Cdc2(as)-M17-GBP-mCherry activity) were added to the culture one hour before release from the G1 arrest. Seven hours after release from the G1 arrest, cells accumulated in G2, and 1-NmPP1 was washed out to remove the inhibition of Cdc2(as)-M17-GBP-mCherry. Mitotic entry was tracked by DNA and septum staining. **b** Representative images of the indicated strains 6 hours after release from the G1 arrest (7 hours after tetracycline addition). Fluorescence images are maximum projections, with the same pixel range applied to all images of the same fluorescent channel. Scale bar, 10 μm. **c** Example maximum-projection images showing DAPI and calcofluor staining of the DNA and septum respectively, taken from the indicated time points after 1-NmPP1 wash-out. Cells were heat-fixed before staining. The displayed pixel range is the same for all images of the same strain, but not between strains; pixel range is chosen to best represent DNA and septum staining. Scale bars, 10 μm. **d** Cells were heat fixed at the indicated time points after 1-NmPP1 wash-out and stained for DAPI and calcofluor to score for passage through mitosis. ‘Aberrant’ includes multiseptate and cut cells. n ≥ 90 cells per time point.

The SPB-tethered strain underwent mitosis with peaks of binucleates of 30-50% (Fig. 2c,d, Supplementary Fig. 2f). There was also an accumulation of aberrant nuclear divisions in later time points indicative of cells that had entered mitosis but encountered difficulties in mitotic progression (Fig 2c,d). This could be due to the artificial SPB tethering system failing to precisely recapitulate normal SPB localisation, or due to the lack of cyclin-substrate docking. In contrast, the non-tethered control strain underwent a clearly lower level of mitosis with reduced peaks of binucleates at 10-20% (Fig. 2c,d, Supplementary Fig. 2f). This was higher than cells driven by Cdc13^HPM^ which did not undergo any mitosis (Fig. 1d), possibly due to increased levels of Cdc13^HPM^ as a consequence of the G2 arrest, and of Cdc2 given there are two gene copies in the strain. We conclude that artificially restoring the localisation of Cdc2-Cdc13^HPM^ at the SPB significantly increases the number of cells entering and proceeding through mitosis.

### Artificial restoration of Cdc2 SPB localisation and CDK substrate phosphorylation

We reasoned that restoration of Cdc2 SPB localisation may rescue mitotic entry due to enhanced CDK substrate phosphorylation. We have previously reported that Cdc13^HPM^ is less efficient than Cdc13^WT^ at bringing about CDK substrate phosphorylation, with half of mitotic CDK substrates phosphorylated less efficiently by Cdc13^HPM^ than Cdc13^WT^ ^7^. Phosphorylation of substrates localised at both the SPB and within the cytoplasm were found to be reduced^7^. We asked if artificial restoration of Cdc2 SPB localisation rescues substrate phosphorylation at the SPB and in the cytoplasm by repeating the experiment in Fig. 2 and taking samples for quantitative phosphoproteomics after 1-NmPP1 removal. We identified 217 previously defined CDK phosphorylation^44^ events in our dataset (Supplementary Table 1). Of these, 51 phosphorylation events were previously classified as HP-sensitive (sites phosphorylated less by Cdc13^HPM^ than Cdc13^WT^), and 50 phosphorylation events were HP-insensitive (sites phosphorylated to wild-type levels by Cdc13^HPM^)^7^. We confirmed that CDK substrate phosphorylation was comparable between both strains before 1-NmPP1 removal in our experiment (Supplementary Fig. 3a).

To identify sites that were increased in phosphorylation as a result of restoring Cdc2 SPB localisation, we identified the ratio of the maximum phosphorylation reached by the SPB-tethered strain and the control strain (defined as the maximum phosphorylation ratio) for each HP-(in)sensitive site (Fig. 3a,b). This experiment detected differences in substrate phosphorylation between the control and the SPB-tethered strains (Fig 3a,b). Phosphorylation events with maximum phosphorylation at least 1.2 times higher in the SPB-tethered than the control strain (which ranged from 1.2 – 1.4 times higher) were predominantly HP-sensitive (Fig. 3c-d). As a group, the HP-sensitive sites also had a higher maximum phosphorylation ratio than HP-insensitive sites (Fig. 3c). Together these results indicate that the sites that increase in response to tethering Cdc2(as)-M17-GBP-mCherry to Cut12-NEGFP are enriched for HP-sensitive sites.

**Figure 3:**
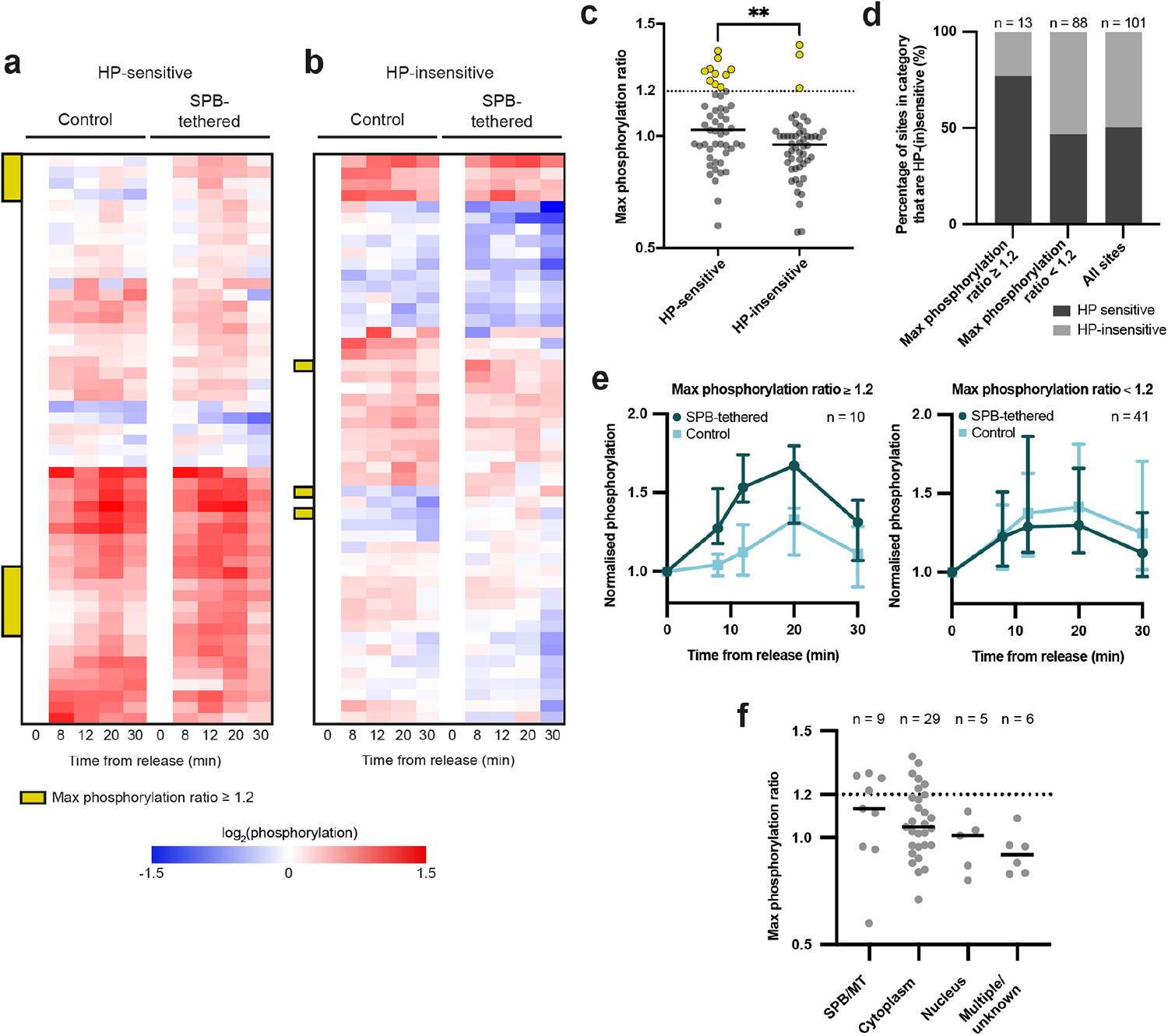
Artificially restoring Cdc13^HPM^ SPB localisation partially rescues CDK substrate phosphorylation at the SPB and cytoplasm. **a, b** Heatmaps showing the log_2_(phosphorylation) of HP-sensitive (**a**) and HP-insensitive (**b**) sites across the time-course for each strain (SPB-tethered and control), normalised to their respective T0. Sites are shown hierarchically clustered. The sites that reach a maximum phosphorylation level at least 20% higher in the SPB-tethered strain than the control (resulting in a max phosphorylation ratio of ≥ 1.2) are indicated by the yellow bar. HP-sensitive n = 51, HP-insensitive n = 50. **c** The maximum phosphorylation ratio of HP-sensitive and HP-insensitive sites. Dashed line indicates a maximum phosphorylation ratio of 1.2, and sites above this are shown in yellow. Solid bars indicate the median of the group. Mann-Whitney rank comparison performed to compare the entire groups; p = 0.0052 (indicated by **). **d** The percentage of sites in each category (max phosphorylation ratio ≥ 1.2, max phosphorylation ratio < 1.2, or all sites) that are either HP-sensitive or insensitive. **e** Median of the normalised phosphorylation of HP-sensitive sites separated according to their max phosphorylation ratio (left: max phosphorylation ratio ≥ 1.2, right: max phosphorylation ratio < 1.2). Error bars show 95% confidence intervals. **f** The max phosphorylation ratio of HP-sensitive sites split according to their reported localisation^7^. Dashed line indicates a max phosphorylation ratio of 1.2, and solid bars indicate the median of the group. Sites with reported localisation to multiple compartments or unknown localisation are in the multiple/unknown category.

Approximately 20% of HP-sensitive sites had a maximum phosphorylation ratio ≥ 1.2, indicating a partial rescue of phosphorylation (Fig. 3e). Comparing the maximum phosphorylation ratio of phosphorylation events in different subcellular compartments showed that the maximum phosphorylation ratio increased in order from the nucleus which was the lowest, increasing in the cytoplasm, to the SPB/microtubules (MTs) which was the highest (Fig. 3f), indicating that some sites with compromised phosphorylation by Cdc13^HPM^ were partially rescued. The phosphorylation events with a maximum phosphorylation ratio ≥ 1.2 included some, but not all, sites from both the SPB/MTs and the cytoplasm (Fig. 3f, Supplementary Fig. 3b). This suggests that tethering Cdc2(as)-M17-GBP-mCherry to Cut12-NEGFP promoted a partial rescue of CDK substrate phosphorylation at both the SPB and within the cytoplasm. We also investigated CDK phosphorylation sites in key G2/M regulators, extending our analysis to include sites that were not in our previous dataset defining HP-sensitivity, finding that some had increased phosphorylation in the SPB-tethered strain (Supplementary Fig. 3c). These included Plo1, which has been shown to interact with Cut12^22^, and had significantly increased phosphorylation on S370 in response to tethering Cdc2(as)-M17-GBP-mCherry to Cut12-NEGFP (Supplementary Fig. 3d). We conclude that artificial tethering of Cdc2-Cdc13^HPM^ to the SPB increases phosphorylation of some CDK substrates at both the SPB and within the cytoplasm.

### Cig2^HPM^ can drive S-phase but has reduced SPB localisation

We next asked if the involvement of the hydrophobic patch in centrosomal localisation, and its importance in cyclin function, is conserved in the S-phase cyclin Cig2. The hydrophobic patch of S- and M-phase budding yeast cyclins have been reported to have different effects on substrate phosphorylation *in vitro*, with the S-phase cyclin hydrophobic patch enhancing phosphorylation of early substrates involved in DNA replication^3^. Therefore, we first asked if Cig2^HPM^ (M168A, L172A, W175A) is able to drive DNA replication, as a critical cell cycle event that relies on CDK phosphorylation of substrates early in the cell cycle. We released *nmt41:cdc13 cig1Δ* cells from a G1 arrest with *cdc13* expression repressed and with either *cig2^WT^*, *cig2^HPM^*, or *cig2Δ* at the endogenous locus and followed S-phase progression using flow cytometry. The populations expressing Cig2^WT^ and Cig2^HPM^ both underwent bulk DNA replication, with a small delay in cells with Cig2^HPM^ compared to Cig2^WT^ (Fig. 4a,b). We conclude that an intact hydrophobic patch is not essential for Cig2 to drive S-phase.

**Figure 4:**
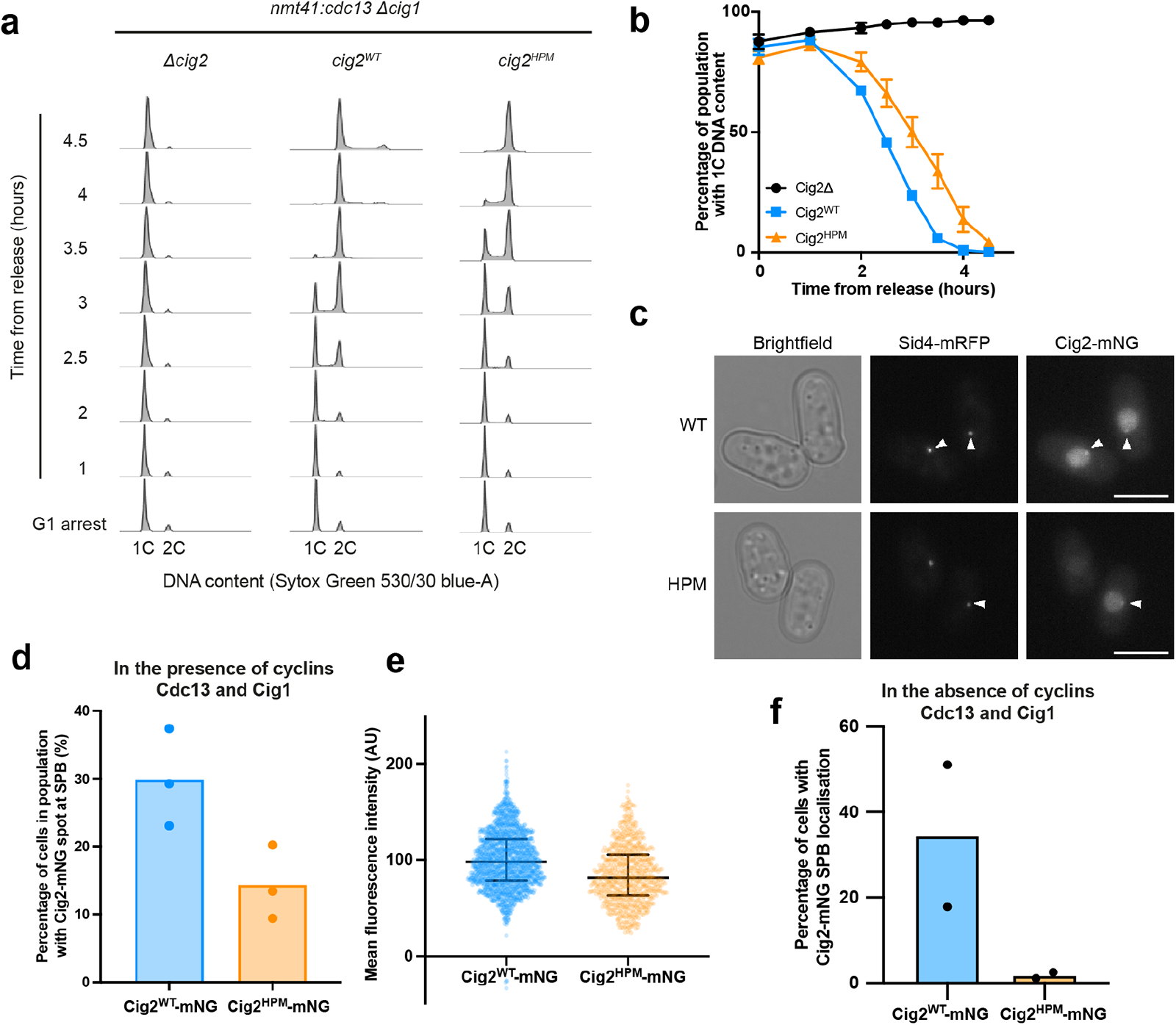
Cig2HPM can drive DNA replication, but is not detected at the SPB in the absence of other cyclins. **a,b** Populations of cells with the cyclins *cdc13* and *cig1* repressed and deleted respectively (*nmt41:cdc13 cig1Δ*) and either *cig1^WT^*, *cig2^HPM^*, or no copy (*cig2Δ*) at the endogenous *cig2* locus were followed after release from a G1 arrest. **a** DNA content of one of three replicates. **b** The percentage of cells with 1C DNA content; mean and SD of three replicates is shown for all time points apart from the *cig2^WT^* strain at 1 and 2 hours, where there are two replicate values. **c-e** An asynchronous population of cells expressing either *cig2^WT^-mNG* or *cig2^HPM^-mNG* at the endogenous *cig2* locus as well as *sid4-mRFP* to mark the SPB were imaged. **c** Representative images of cells with Cig2^WT^-mNG and Cig2^HPM^-mNG at the SPB (marked by arrowheads). Scale bars, 5 μm. **d** Percentage of cells in the population with Cig2^WT^-mNG and Cig2^HPM^-mNG at the SPB; bar shows median of three replicates (individual replicates are represented by dots). **e** Mean cellular fluorescence intensity of Cig2^WT^-mNG and Cig2^HPM^-mNG; dots represent individual cells and bars represent the median and IQR of the population. One replicate of three is shown, and all values are shown after the mean autofluorescence intensity has been subtracted. Cig2^WT^-mNG n = 1658 and Cig2^HPM^-mNG n = 746 cells. **f** Cells with the cyclins *cdc13* and *cig1* repressed and deleted respectively, as well as *sid4-mRFP* and either *cig2^WT^-mNG* or *cig2^HPM^-mNG* at the endogenous *cig2* locus were released from a G1 arrest and followed with time-lapse imaging from 1.5 hours after release for a further 4.5 hours. Graph shows percentage of cells with Cig2-mNG at the SPB at any point during the time-lapse. Bar shows median of two replicates; individual replicates are shown by dots.

We also investigated Cig2^HPM^ SPB localisation. Endogenous Cig2^WT^-mNG and Cig2^HPM^-mNG were both detected at the SPB in relatively short cells (indicating they were early in the cell cycle) in an asynchronous population, suggesting that, like Cdc13^7^, there is a HP-independent mechanism of Cig2-mNG SPB localisation (Fig. 4c,d and Supplementary Fig. 4a). Roughly twice as many cells had detectable Cig2^WT^-mNG SPB foci than Cig2^HPM^-mNG (Fig. 4d), so it is possible that as with Cdc13, the hydrophobic patch of Cig2 is also involved in its localisation to the SPB. However, Cig2^HPM^-mNG is expressed at a lower level in the population, at roughly 80% of the Cig2^WT^-mNG level (Fig. 4e), which may contribute to a reduction in detectable Cig2^HPM^-mNG SPB foci.

The HP-independent mechanism of Cdc13 SPB localisation is not detected in the absence of wild-type Cdc13, Cig1 and Cig2 (Fig. 1e,f). We therefore asked whether the same was true for Cig2. We released *Δcig1* cells from a G1 arrest whilst repressing *cdc13* transcription and monitored endogenous Cig2-mNG localisation by timelapse microscopy. There was variability in both the timing and the level of peak Cig2^WT^-mNG and Cig2^HPM^-mNG after release (Supplementary Fig. 4d,e). Cig2^HPM^-mNG peaked slightly later and lower than Cig2^WT^-mNG, consistent with our data that Cig2^HPM^-mNG is expressed at a lower level in an asynchronous population (Supplementary Fig. 4d,e). We altered our spot detection workflow (Supplementary Fig. 1) to take account of the Sid4-mRFP signal being too weak to mask the SPB in all time points of the timelapse (see Methods). The assay did not detect Cig2^WT^-mNG SPB localisation in all cells, however there were almost no cells with Cig2^HPM^-mNG SPB localisation in contrast to Cig2^WT^-mNG (Figure 4f). We confirmed that the decrease in detectable SPB foci is not due to the lower expression of Cig2^HPM^-mNG, as restricting the analysis to cells of similar expression levels did not affect the result (Supplementary Fig. 4f,g). Thus, in the absence of Cdc13 and Cig1, we did not detect Cig2^HPM^-mNG at the SPB. This suggests that the hydrophobic patch of Cig2 mediates its localisation to the SPB, and that the hydrophobic patch-independent mechanism of Cig2-mNG SPB localisation involves wild-type Cig2, Cdc13 and/or Cig1.

## Discussion

We have investigated the role that cyclin-CDK localisation at the centrosome plays in ensuring proper progression through the eukaryotic cell cycle. This work has taken advantage of the finding that mutation of the hydrophobic patch of the mitotic cyclin Cdc13 in the fission yeast *S. pombe* disrupts the localisation of the Cdc13-Cdc2 cyclin-CDK at the SPB (centrosome) during G2 of the cycle^7^. The importance of this centrosomal localisation on mitotic entry and on the phosphorylation of Cdc13-Cdc2 substrates, as well as the role of the S-phase Cig2-Cdc2 cyclin-CDK hydrophobic patch, have been investigated.

Cells driven by the hydrophobic patch mutant Cdc13^HPM^-sfGFP arrest in G2 and do not undergo mitosis. In this experiment there is no detectable Cdc13^HPM^-sfGFP enrichment at the SPB. Artificially restoring Cdc13^HPM^ localisation at the SPB promotes a significant increase in mitotic entry although there were some aberrant nuclear divisions. It also rescues CDK phosphorylation of some substrates located both at the SPB and in the cytoplasm. Artificial re-localisation to the SPB by tethering to the SPB component Cut12 can be expected to provide increased cyclin-CDK activity in the vicinity of Cut12, directly aiding phosphorylation of SPB substrates. Our results also indicate that the Cdc2-Cdc13^HPM^ tethering at the SPB increases the phosphorylation of some CDK substrates within the cytoplasm. This could be due to increased CDK activity in the cytoplasm brought about by activation of the CDK at the SPB by positive feedback mechanisms^24,35^ and subsequent propagation of this activity to the cytoplasm. Alternatively, some CDK substrates may be phosphorylated at the SPB and subsequently diffuse to the cytoplasm. Either way, these results support the view that the SPB is a critical hub for controlling mitosis^12,24^. Given that re-localisation to the SPB does not increase phosphorylation of all CDK substrates, our results suggest that the cyclin hydrophobic patch is also important for substrate docking *in vivo* as has been previously proposed^2,3,36,45^. We therefore suggest that our *in vivo* studies support roles for the mitotic cyclin hydrophobic patch for both centrosomal localisation and for substrate docking to enhance CDK substrate phosphorylation, and also emphasise the importance of G2 CDK centrosomal localisation for the subsequent entry into mitosis. Concentration of CDK regulators possibly working through feedback loops, or CDK substrates being co-located with cyclin-CDK, could promote the cell cycle transition into and through mitosis.

Cyclin-substrate docking of S-phase cyclins has been shown to enhance *in vitro* phosphorylation of CDK substrates involved in DNA replication in budding yeast, suggesting that the hydrophobic patch may be important for the early cell cycle event of S-phase^3^. However, in fission yeast the mitotic cyclin Cdc13^HPM^ is able to drive S-phase^7^. We therefore tested if the role of the hydrophobic patch in centrosomal localisation is conserved in the *S. pombe* S-phase cyclin Cig2, and the importance of the Cig2 hydrophobic patch for S-phase. Disruption of the hydrophobic patch of Cig2 revealed both hydrophobic patch-dependent and independent mechanisms of Cig2 SPB localisation, similar to what is found for the mitotic cyclin Cdc13^7^. Also like Cdc13^HPM^ ^7^, we found that Cig2^HPM^ can drive bulk DNA replication, indicating that the Cig2 hydrophobic patch is not essential to bring about S-phase. These results establish the importance of the hydrophobic patch in centrosomal localisation of both S-phase and M-phase cyclins in fission yeast, as has also been observed in human cyclins^5–7^.

Our results strengthen the view that the centrosomal localisation of cyclin-CDK plays an important role for cell cycle regulation. We have shown that phosphorylation of some CDK substrates is enhanced by cyclin-CDK centrosomal localisation, and without this localisation substrate phosphorylation is insufficient to bring about mitosis. The enhancement of substrate phosphorylation may be due to activation of the CDK brought about by co-location of CDK and CDK regulators, or due to the co-location of CDK and its substrates. Whilst centrosomes are not essential for mitosis in *Drosophila* or in some vertebrate cell lines^12,17–19^, it is possible that in the absence of a centrosome, co-location of CDK and its regulators or substrates elsewhere in the cell may be sufficient to bring about mitotic entry^13^. Perhaps what matters is the co-location of components critical for mitotic entry signalling, rather than precisely where that co-location occurs. The localisation of a number of core cell cycle regulators including cyclin-CDK at the metazoan centrosome^9,46,47^, and the conservation of the role of the Cyclin B hydrophobic patch in centrosomal localisation^5,7^, suggest that our findings are relevant for mitotic control throughout eukaryotes.

## Methods

### Strain construction and growth conditions

Fission yeast media and cell culture techniques are described previously^48^. Cells were grown in Edinburgh minimal media (EMM) supplemented with adenine, leucine and/or uridine at 0.15 g/L where required. Cells were grown at 32 °C apart from when imaging Cig2-mNG in an asynchronous population (Fig. 4c-e), for which cells were grown at 25 °C. G1 arrests by nitrogen starvation (detailed below), were also performed at 25 °C. Strains PN6033 and PN6034, which include the tetracycline-inducible *tetP-cdc2(as)-M17-GBP-mCherry*, were grown in media supplemented with 10 μM 1-NmPP1 before the experiment to inhibit activity of any Cdc2(as)-M17-GP-mCherry protein produced by potential leaky expression. Strain construction was performed by mating and subsequent random spore analysis, or by transformation^48^ and confirmed by colony PCR. The following plasmids were both gifts from Michael Nick Boddy and bought from Addgene: pFA6a-KanMX-tetO-pCyc1 (TetP) (Addgene plasmid # 41023; http://n2t.net/addgene:41023; RRID:Addgene_41023); and pDM291-tetR-tup11Δ70 (Addgene plasmid # 41027; http://n2t.net/addgene:41027; RRID:Addgene_41027)^49^. pFA6a-GBP-mCherry-kanMX6 was a gift from Quanwen Jin (Addgene plasmid # 89068; http://n2t.net/addgene:89068; RRID:Addgene_89068)^50^. The strains used in this study and the related plasmids used to construct them are listed in Supplementary Table 2.

### Cell cycle arrest and release

G1 arrest of cells was performed by taking an exponentially growing culture at a density of 2 x 10^6^ cells/mL and washing at least three times in media containing no ammonium chloride and no supplements. Cells were held in the nitrogen starvation for between 16-19 hours at 25 °C, before release by washing into EMM media which contains ammonium chloride with the relevant supplements. To repress the thiamine-repressible *nmt41* promoter, thiamine hydrochloride (Sigma) dissolved in water was added to a final concentration of 30 µM one hour before release from the nitrogen arrest and to the media used to release the cells. To induce protein from the *TetP* promoter described previously^49^, anhydrotetracycline hydrochloride (Sigma) dissolved in DMSO was added to a final concentration of 1.25 µg/mL.

### Fluorescence microscopy and image analysis

Fluorescence microscopy was performed with a Nikon Ti2 inverted microscope, equipped with a Perfect Focus System and Okolab environmental chamber, and a Prime sCMOS or BSI camera (Photometrics). Micro-Manager v2.0 software (Open-imaging) was used to control the microscope^51^. Fluorescence excitation was carried out using a SpectraX LED light engine (Lumencor) fitted with the following standard filters: 395/25 for imaging DAPI/calcofluor, 470/24 for imaging sfGFP/eGFP; and 575/25 for imaging mRFP; with either a dual-edge ET-eGFP/mCherry dichroic beamsplitter (Chroma 59022bs), or a 409/493/573/652 nm BrightLine® quad-edge dichroic beamsplitter (Semrock). The following emission filters were used: Semrock, 438/24 nm BrightLine® single-band bandpass filter for imaging DAPI/calcofluor; Chroma, ET – EGFP single-band bandpass filter ET525_50m for imaging sfGFP/eGFP; and Semrock, 641/75 nm BrightLine® single-band bandpass filter FF02_641_75 for imaging mCherry/mRFP. A 100X Plan Apochromat oil-immersion objective (NA 1.45) was used and microscopy performed at 25 °C for still imaging, or 32 °C for time-lapse.

Time-lapse imaging was performed on agar pads, made by dissolving agarose to a final concentration of 2% w/v in EMM media supplemented with leucine, adenine and thiamine. The solution was used to fill a gene frame (Thermo Fisher) and allowed to solidify. Cells were mounted in their growth media on the agar pads, and imaged. ImageJ software (NIH) was used to measure pixel intensity, adjust brightness and contrast, apply a pixel range to images, and render maximum projection images^52^. Where images are shown, the same pixel range has been applied to all images of the same channel within a figure panel, unless otherwise stated.

For analysis of still imaging, cells were segmented using Ilastik^53^. For the whole-cell intensity measurements of Cig2-mNG in an asynchronous population (Fig. 4e), background value of autofluorescence was subtracted from the measurements. To find the background value of autofluorescence, the mean cellular fluorescence intensity was measured of a population of cells with the same genetic background but without Cig2-mNG, and the median of this population was taken as the value of autofluorescence.

For analysis of time-lapse images, for faster segmentation, cells were segmented and labelled using a custom script in Matlab 9.13.0 R2022b (Mathworks). To identify edges of cells more clearly, cells were segmented on images 1 μm below the focal plane. First, cells were segmented using the “imbinarize” function with an adaptive threshold algorithm. Any regions of the out-of-cell background were removed using an intensity threshold as cells contained higher intensity values. Any gaps formed inside cells were filled using the “imfill” function. The binary masks of cells were eroded to facilitate the detection of cell boundaries. A watershed algorithm was used using the “watershed” function to identify cells and their boundaries. The resulting masks were dilated to restore their normal size and the watershed algorithm was used again to check if cell boundaries were accurately detected. Holes in the binary masks were filled using the “imfill” function. Any subsequent out-of-cell artifacts were removed using size and intensity thresholds. The masks of cells were labelled with numbers using the “labelmatrix” function for tracking analysis. Labelled cells were then tracked using the Lineage Mapper plugin in FIJI^54^. Whole-cell intensity measurements were performed by taking the mean fluorescence intensity in the cell mask.

Spot detection to identify Cdc13 and Cig2 localisation to the SPB was performed using the trackmate v7.10.0 FIJI plugin^55,56^. A DoG detector was used for all experiments with the estimated spot diameter set to 0.35 μm, and the following parameters selected: pre-process with median filter; sub-pixel localisation. The parameters used to filter spots were determined empirically for each different experiment, with the same settings being used between all strains and all replicates of any given experiment. Due to Sid4-mRFP signal being close to background in the time-lapse, Sid4-mRFP spots were not detected in all frames of the timelapse; to account for this the spot detection workflow outlined in Supplementary Fig. 1 was adapted. Cells were counted as having Cig2-mNG at the SPB if they had either a Cig2-mNG spot that intersected with a Sid4-mRFP spot in any given frame of the timelapse, or a Cig2-mNG spot in two out of three consecutive frames with no detected Sid4-mRFP spot.

### Flow cytometry and cell cycle progression analysis

DNA content analysis was performed using flow cytometry. Cells were fixed in 70% ice-cold ethanol, before being washed and resuspended in 50 mM sodium citrate. Samples were incubated with 0.1 mg/mL RNase A at 37 °C for at least 3 hours. DNA was stained with Sytox Green (1 µM, Invitrogen). 100,000 events were acquired on either an LSRII or BD LSRFortessa, using FACS Diva software. DNA content is displayed on a linear scale after gating for single cells in FlowJo v10.7.2 (Treestar Inc). Analysis of the percentage of cells with 1C DNA content in Fig. 4b was performed by gating around the 1C peak in the histograms represented in Fig. 4a using FlowJo v10.3.

Binucleation analyses and detection of aberrant divisions were performed by heat-fixing cells on a microscope slide at 70 °C before staining with 4’,6-diamidino-2-phenylindole (DAPI) (SlowFade Diamond Antifade Mountant with DAPI, Invitrogen) and Calcofluor (Sigma). Cells that displayed the ‘*cut*’ phenotype (a septum intersecting an undivided nucleus, or two foci of DAPI staining very close to a septum) were included in the ‘aberrant’ category, as were cells that had a single nucleus on one side of a septum, or multiple septa.

### Protein extraction and phosphoproteomics

Protein extracts were prepared by addition of ice-cold 100% w/v trichloroacetic acid (TCA) to a final volume of 10% to a cell culture. Cells were kept on ice for at least 30 minutes before being washed in ice-cold acetone. Dry pellets were stored at -80 °C, before being washed and resuspended in lysis buffer (8 M urea, 50 mM ammonium bicarbonate, 5 mM ethylenediamine tetraacetic acid (EDTA), 1 mM phenylmethylsufonyl fluoride (PMSF), 1X protease inhibitor cocktail set III (EDTA free, CalBiochem), and 1X PhosSTOP phosphatase inhibitor cocktail (Roche)). Cells were lysed by two rounds of beating with glass beads at 4 °C using a FastPrep120 at 5.5 m/s for 45 s, followed by one round of beating at 6.5 m/s for 40 s, with the samples placed on ice for 2 min between rounds of beating. The supernatant was collected after cell debris was pelleted at 13,000 rpm in a table-top microcentrifuge at 4 °C.

For phosphoproteomic analysis, lysates (200 μg) were digested using SP3 beads according to manufacturer instructions using Trypsin and LysC. Following digestions samples were labelled with TMT10plex Isobaric Label Reagents kit (Thermo Fisher) as previously described^57^, with the exception that peptides were desalted using a desalting column (89852, Thermo Fisher). Samples were normalised using label amounts as previously described^58^ and subjected to sequential TiO2 and Fe-NTA phosphopeptide enrichment protocols according to manufacturer instructions. Resulting phosphopeptides and whole cell lysates (flow through) were fractionated using Pierce High pH Reverse-Phase Peptide Fractionation kit (Thermo Fisher). Samples were dried and reconstituted in 0.1% trifluoroacetic acid (TFA). Peptides were loaded onto trap column and separated using Easy Spray 50cm (Ultimate3000) using 180 min gradient (mobile phase A 0.1% TFA – 95%, 5% DMSO, mobile phase B -75 %ACN, 5% DMSO, 5% water). For the gradient buffer B concentration was 2% – 0min, 8%-5.5min, 40%-153min, 95%-155min, 2%-165min. Data were acquired using Orbitrap Eclipse using SPS-MS3 method with the following settings: for MS1 Orbitrap resolution was set to 120000, AGC target to standard and automatic injection time, for MS2 Orbitrap resolution was set at 30000 with fragmentation CID 35, and standard AGC target and injection time, and for MS3 Orbitrap resolution was set to 50000, HCD fragmentation was set at 65 and AGC target was set to 200% with 60ms maximum injection time.

### Phosphoproteomic data analysis

The dataset was searched on MaxQuant^59^ v1.6.14 against *Schizosaccharomyces pombe* proteome FASTA file extracted from UniProt^60^, amended to include common contaminants and the introduced exogenous Cdc2(as)-M17-GBP-mCherry construct. The default MaxQuant parameters were used with Phospho(STY) added as a variable modification. Subsequent analysis was performed in Perseus v1.6.14.0^61^. Different multiplicities of the same site are treated as separate phosphorylation events. The data was filtered to remove contaminant and reverse hits, and to retain only sites that appeared in all time-points of the time-course and had a localisation probability of at least 0.7. All phosphorylation events were normalised to the median value of their time-point to account for potential mixing errors. All time-points of all phosphorylation events were then normalised to their respective T0. Sites were further normalised to their respective T0-normalised protein change (sites without proteome data for all time points were excluded). In all line graphs where phosphoproteomic data is presented, the data is presented in this format (protein-normalised, T0-normalised phosphorylation) and this is referred to as the normalised phosphorylation. To display phosphorylation in a heat-map, phosphorylation events were hierarchically clustered using Euclidean distance with complete linkage and without pre-processing with k-means.

To calculate the max phosphorylation ratio of a given phosphorylation event, the maximum normalised phosphorylation reached in the time-course was identified for both strains. The ratio was calculated using: SPB-tethered max phosphorylation/control max phosphorylation. Substrate localisation information was compiled from published literature in our previous study^7^.

### Data representation

Statistical tests performed are detailed in the figure legends and were performed using GraphPad Prism 9. Normality was checked by the D’Agostino and Pearson normality test, and Mann-Whitney rank comparison was used to compare conditions that were not normally distributed.

## Acknowledgements

We thank Jessica Greenwood and Sarah Willich for critical reading of this manuscript, Jack Mead for help with image analysis scripts, and Scott Curran for providing the Ilastik cell segmentation pipeline. We thank all members of the Nurse laboratory for helpful discussions. We are grateful to Iain Hagan, Michael Nicholas Boddy, Silke Hauf, Quanwen Jin and Masamitsu Sato for providing strains and plasmids. We also thank the Flow Cytometry and Light Microscopy facilities at the Francis Crick Institute for their assistance. This work was supported by the Francis Crick Institute which receives its core funding from Cancer Research UK (CC2003), the UK Medical Research Council (CC2003), and the Wellcome Trust (CC2003). In addition, this work was supported by the Wellcome Trust Grant to PN (grant number 214183 and 093917), The Lord Leonard and Lady Estelle Wolfson Foundation, Woosnam Foundation and Breast Cancer Research Foundation (BCRF-22-117). N.K was supported by an EMBO postdoctoral fellowship (grant number EMBO ALTF 705-2020) and a HFSP postdoctoral fellowship (grant number HFSP LT000587/2021-L). For the purpose of Open Access, the author has applied a CC BY public copyright licence to any Author Accepted Manuscript version arising from this submission.

## Author contributions

E.L.R, S.B. and P.N. conceptualised the study. P.N. supervised the study. E.L.R conducted experiments and analysed data. N.K. developed custom Matlab scripts for image processing. T.A. performed preparation of samples for mass spectrometry and data acquisition. E.L.R and P.N wrote the manuscript with input from all authors.

## Competing interests statement

The authors declare no competing interests.

## Notes

### Competing Interest Statement

The authors have declared no competing interest.

