## Supplementary information for "CDK activity at the centrosome regulates the cell cycle"

Supplementary Table 1: Condensed phosphoproteomic dataset.

Supplementary Table 2: Fission yeast strains and plasmids used in this study.

Supplementary video 1: $Cig2^{WT}$-mNG localisation after release from G1 arrest

Supplementary video 2: $Cig2^{HPM}$-mNG localisation after release from G1 arrest

### Supplementary figure 1

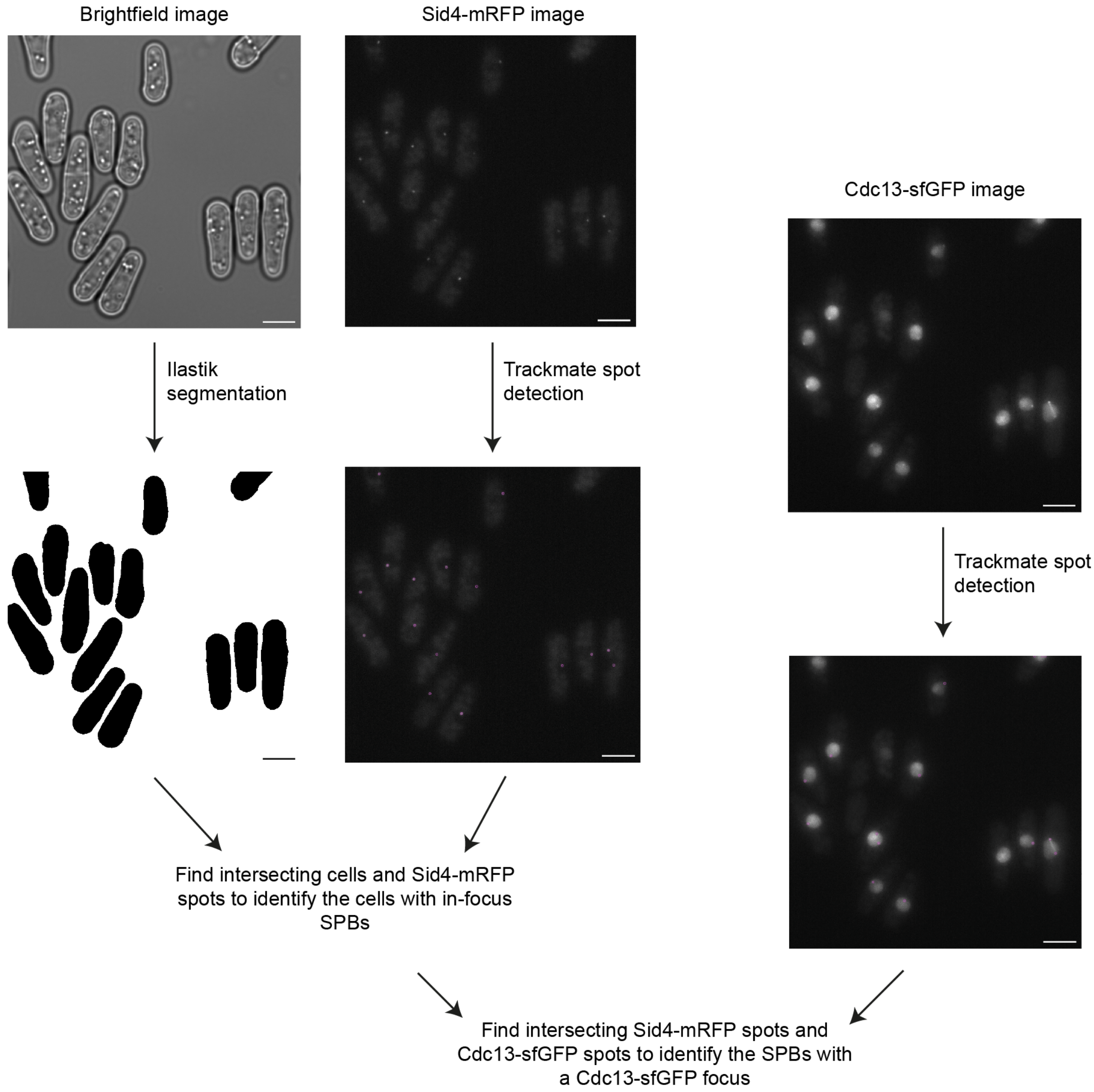


**Supplementary Figure 1: Automated spot detection method to identify Cdc13 SPB localisation**

Brightfield images were segmented using Ilastik to generate cell masks. The trackmate Fiji plugin [1,2] was used to detect Sid4-mRFP and Cdc13-sfGFP spots from maximum projection images (spots are shown overlaid on images in magenta). Scale bars, 5 µm.

### Supplementary Figure 2

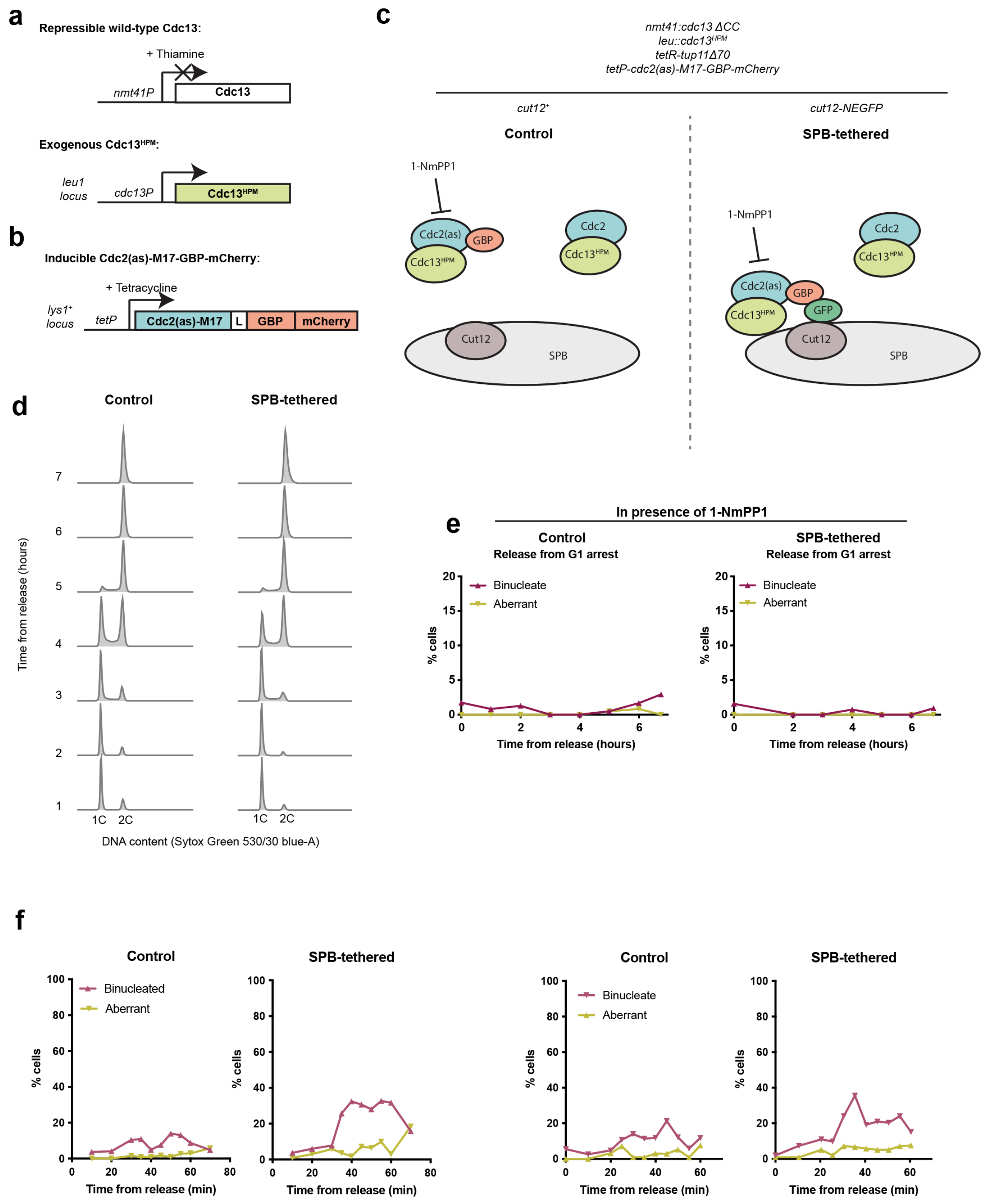


**Supplementary Figure 2: Artificial tethering of Cdc2 to the SPB**

**a** Cdc13 conditional expression system used in the experiment detailed in Figure 2. Wild-type Cdc13 is placed under the thiamine-repressible *nmt41* promoter. Cells also express an exogenous copy of Cdc13$^{HPM}$, expressed under the *cdc13* promoter and inserted into the *leu1* locus. **b** System to conditionally express and activate an exogenous copy of Cdc2. The analogue sensitive allele Cdc2(as)-M17[3] is C-terminally tagged with GBP-mCherry, and placed under a tetracycline-inducible promoter (*tetP*)[4]. The addition of tetracycline induces the expression of the construct, and the addition of the ATP analogue 1-NmPP1 inhibits the activity of the construct. **c** System to artificially restore Cdc2 SPB localisation. The Cdc13 conditional expression system outlined in (a) is used to ensure that upon addition of thiamine, the only copy of Cdc13 that is expressed is $Cdc13^{HPM}$. The G1/S cyclins *cig1* and *cig2* are also genetically deleted (referred to as *ΔCC*). In addition to the endogenous Cdc2, the system outlined in (b) is used to conditionally express and activate an exogenous copy of Cdc2(as)-M17-GBP-mCherry. In the strain containing Cut12-NEGFP (SPB-tethered strain), the exogenous Cdc2(as)-M17-GBP-mCherry is tethered to the SPB; in the strain containing wild-type Cut12 (control strain) it is not tethered to the SPB. **d** DNA content of cells after release from the G1 arrest, measured by flow cytometry. **e** Cells were heat-fixed at the indicated time-points after release from the G1 arrest, and scored for binucleation and septation to measure progression through mitosis. n > 100 cells per time point. **f** Two further replicates of the data shown in Fig. 2d. n > 100 cells per time point.

### Supplementary figure 3

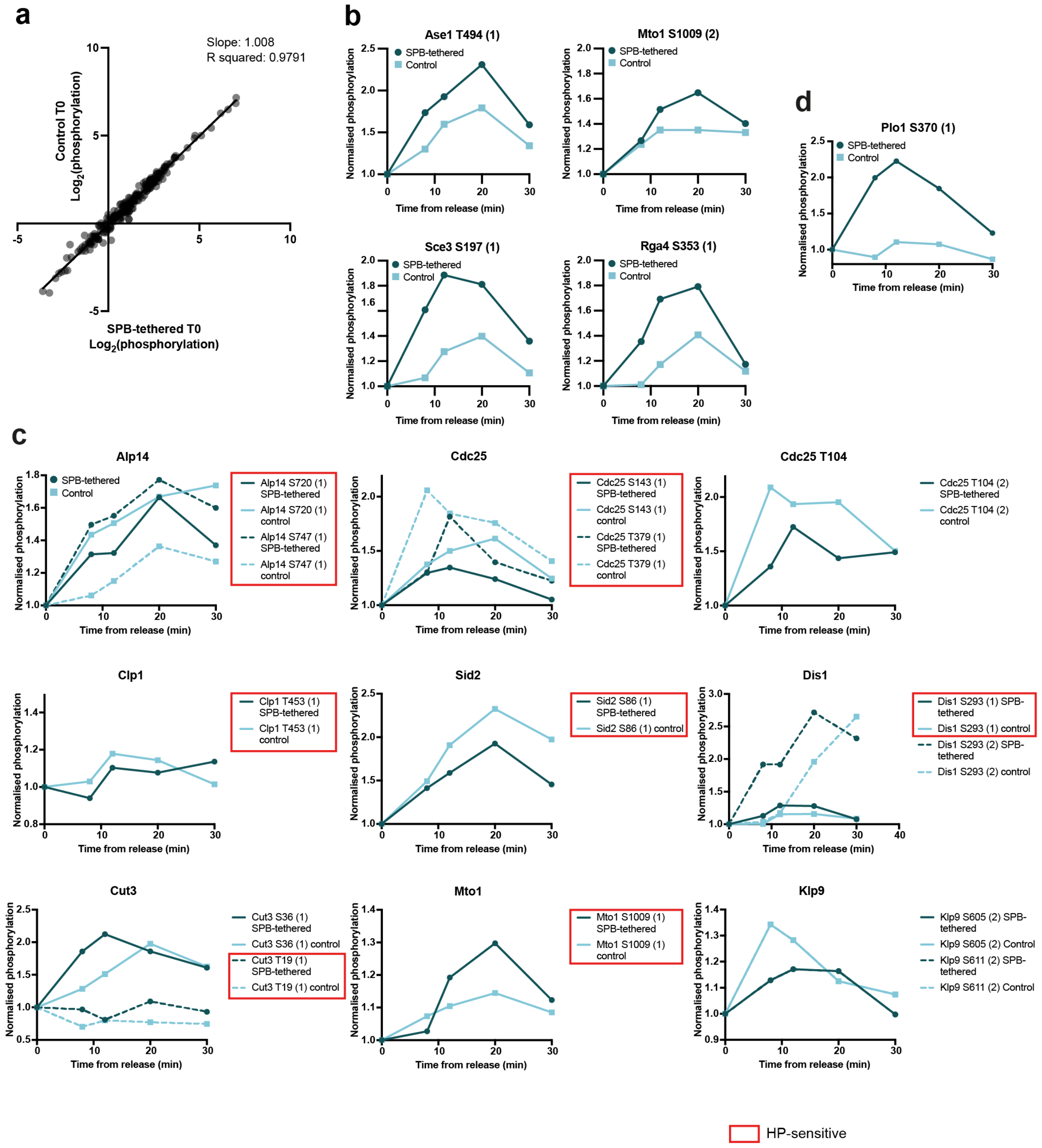


**Supplementary Figure 3: CDK substrate phosphorylation after tethering Cdc2 to the SPB**

**a** $Log_2$(phosphorylation) of the T0 samples of previously defined CDK sites[5] before normalisation. Line represents linear regression. n = 224 phosphorylation events. **b** Normalised phosphorylation of example HP-sensitive sites on proteins reported to localise to the SPB (top two graphs) and cytoplasm (lower two graphs) that have a max phosphorylation ratio ≥ 1.2. Phosphosite position and multiplicity are both indicated, for example S1009 (2) refers to serine 1009, multiplicity 2. **c** Normalised phosphorylation of example key G2/M CDK substrates. Phosphorylation events enclosed by a red box have been previously identified as HP-sensitive[6]. Phosphorylation events not enclosed by a box were not in the dataset that defined HP-sensitive sites. The SPB-tethered strain is always shown in dark blue, and the control strain in light blue. Where two phosphosites or multiplicities are plotted on the same graph, one is indicated by a solid line and the other in a dashed line. Phosphorylation multiplicity is indicated in brackets in the legend. Both Klp9 phosphosites displayed the same phosphorylation, so the lines are overlaid. **d** Normalised phosphorylation of Plo1 S370 (multiplicity 1), which has previously been identified as a CDK site[5].

### Supplementary Figure 4

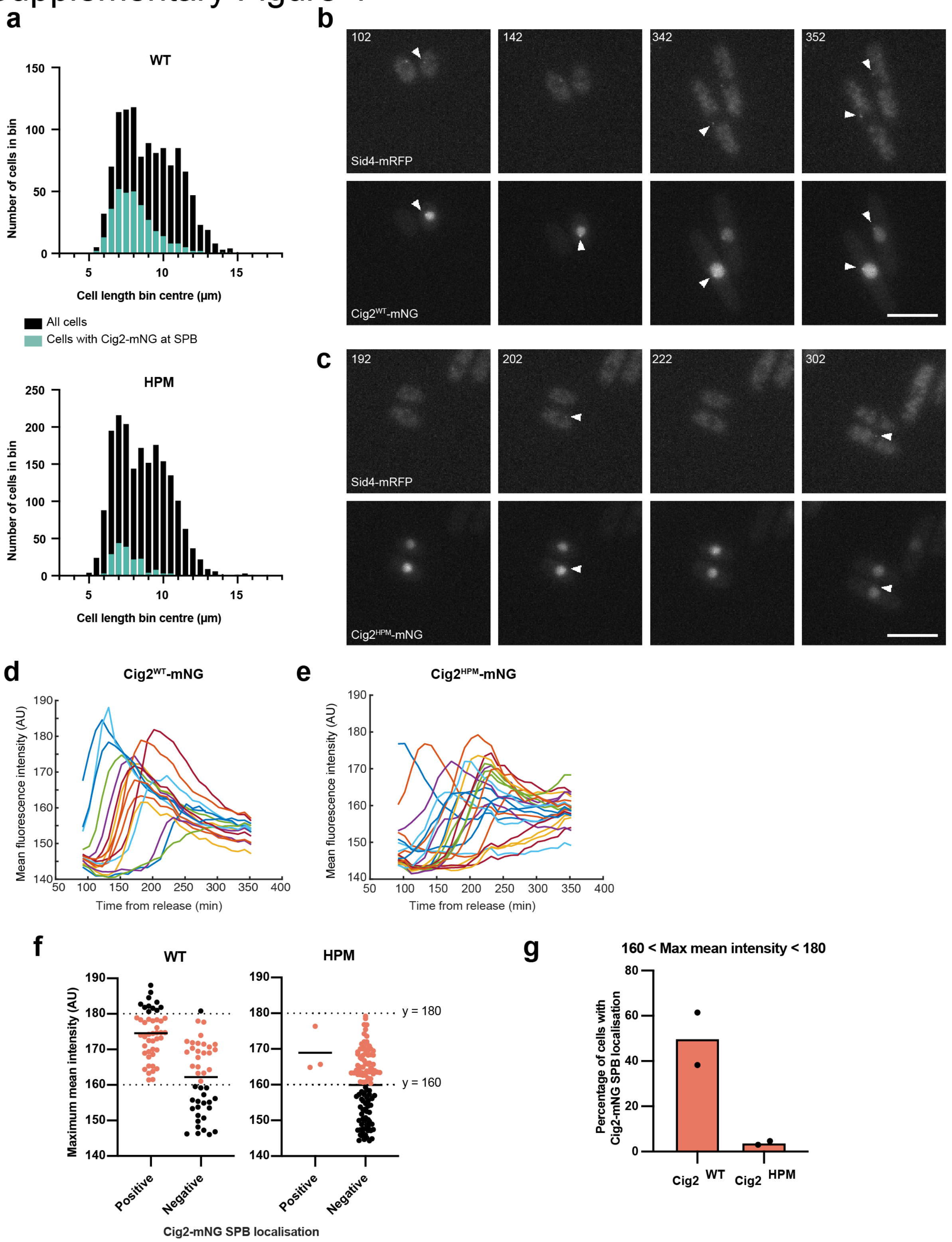


**Supplementary Figure 4: Cig2$^{HPM}$-mNG localises to the SPB only in the presence of Cdc13 and/or Cig1**

**a** Histogram of cell lengths (collected into 0.5 µm bins) of cells with Cig2$^{WT}$-mNG or Cig2$^{HPM}$-mNG at the SPB (green bars) overlaid on of all cells in the population (black bars). One replicate of three is shown. Cig2$^{WT}$-mNG all cells n = 1115, Cig2$^{HPM}$-mNG all cells n = 1909. **b-c** Maximum projection images of example time-points of representative cells expressing Cig2$^{WT}$-mNG (**b**) or Cig2$^{HPM}$-mNG (**c**) after release from a G1 arrest with *cdc13* expression repressed and *cig1Δ*. Sid4-mRFP marks the SPB. Both Cig2$^{WT}$-mNG cells had Cig2$^{WT}$-mNG SPB localisation; arrows indicate Cig2$^{WT}$-mNG foci and corresponding Sid4-mRFP foci where detected. Neither cell expressing Cig2$^{HPM}$-mNG had Cig2$^{HPM}$-mNG at the SPB; arrows indicate location of Sid4-mRFP foci. Time-stamp (top left) represents minutes from release from the G1 arrest; imaging started at 92 minutes. Scale bars, 10 μm. Data from the full time-lapse (including all time points) for these cells is shown in the Supplementary videos 1 and 2 **d-e** Individual cell traces showing mean fluorescence intensity of Cig2$^{WT}$-mNG (**d**) or Cig2$^{HPM}$-mNG (**e**) in cells from one representative field of view from one replicate of two. Cig2$^{WT}$-mNG n = 16, Cig2$^{HPM}$-mNG n = 23. **f** The maximum of the mean cellular fluorescence intensity reached during the time-lapse for each cell; cells are separated by strain and whether they have Cig2-mNG at the SPB. Bars represent median of each population. One replicate of two is shown. Cig2$^{WT}$-mNG positive n = 46, negative n = 44, Cig2$^{HPM}$-mNG positive n = 3, negative n = 114. **g** The percentage of cells with Cig2-mNG at the SPB after restricting analysis to cells with a maximum mean intensity between 160 and 180 (represented in red in **f**) to reduce the impact of expression differences between *cig2$^{WT}$-mNG* and *cig2$^{HPM}$-mNG*. The lower bound of 160 AU is approximately the lowest level at which a Cig2-mNG SPB focus was detected; the higher bound of 180 AU is approximately the maximum level reached by Cig2$^{HPM}$-mNG. Dots represent the two individual replicates and the bar represents the median of these.
