## Supplementary figures and images for "CDK activity at the centrosome regulates the cell cycle"

### Supplementary video 1

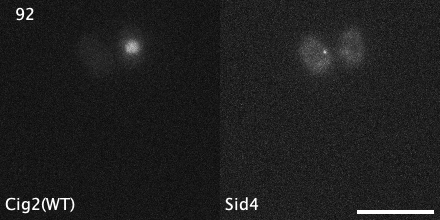

### Supplementary video 2

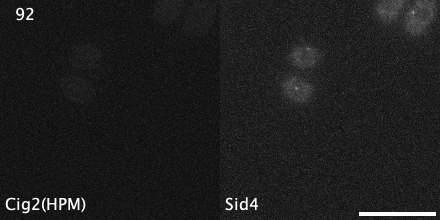
